# Network-based clustering for drug sensitivity prediction in cancer cell lines

**DOI:** 10.1101/764043

**Authors:** Maryam Pouryahya, Jung Hun Oh, James C. Mathews, Zehor Belkhatir, Caroline Moosmüller, Joseph O. Deasy, Allen R. Tannenbaum

**Affiliations:** Department of Medical Physics, Memorial Sloan Kettering Cancer Center, NY, USA; Department of Chemical and Biomolecular Engineering, Johns Hopkins University, Baltimore, USA; Department of Mathematics, University of California, San Diego, La Jolla, USA; Departments of Computer Science and Applied Mathematics & Statistics, Stony Brook University, NY, USA

## Abstract

The study of large-scale pharmacogenomics provides an unprecedented opportunity to develop computational models that can accurately predict large cohorts of cell lines and drugs. In this work, we present a novel method for predicting drug sensitivity in cancer cell lines which considers both cell line genomic features and drug chemical features. Our network-based approach combines the theory of optimal mass transport (OMT) with machine learning techniques. It starts with unsupervised clustering of both cell line and drug data, followed by the prediction of drug sensitivity in the paired cluster of cell lines and drugs. We show that prior clustering of the heterogenous cell lines and structurally diverse drugs significantly improves the accuracy of the prediction. In addition, it facilities the interpretability of the results and identification of molecular biomarkers which are significant for both clustering of the cell lines and predicting the drug response.

## Introduction

In recent years, there have been significant advances in linking pharmacological data with tumor genomics and transcriptomics. This has been driven in part by advances in high-throughput technologies to develop data sets of genomic information of cancer cell lines accompanied with their drug sensitivities. Pioneers of such datasets are the NCI-60 data base [1], Genomics of Drug Sensitivity in Cancer (GDSC) project [2], and the Cancer Cell Line Encyclopedia (CCLE) project [3]. Collectively, these studies have demonstrated that pharmacogenomic profiling of cancer cell lines from clinical tumor samples could help with development of new cancer therapies [4, 5]. The NCI-60 project is one of the first established tools for *in vitro* drug screening and has significantly improved the philosophy and research of human cancer drugs [1, 6]. This panel has led to many important discoveries, including a general advance in the understanding of the mechanism of cancer and the action of drugs [7, 8]. However, this panel only used 60 cell lines, which limits its use for developing predictive models. The Genomics of Drug Sensitivity in Cancer (GDSC) database (http://www.cancerRxgene.org), on which we focus in this study, annotates a comprehensive landscape of drug responses of ∼1000 human cancer cell lines to 265 anti-cancer agents. Importantly, the genomic and transcriptomic profiles of the entire cancer cell lines used in GDSC were extensively characterized as a part of the COSMIC cell line project (CCLP) (https://cancer.sanger.ac.uk/cell_lines). These resources have the potential to link anticancer drug sensitivity to detailed genomic information and facilitate the discovery of molecular biomarkers of drug response. However, powerful analytical tools are needed to cope with the high-dimensionality and complexity of these datasets. In recent years, there are have been a variety of approaches proposed for identification of drug sensitivity in cancer cell lines. Staunton *et al.* [9] first developed a weighted voting classification model for anticancer drug sensitivity on NCI-60 gene expression data. Also, a number of machine learning techniques have been utilized for supervised learning of drug responses. These techniques may be partitioned into regression to predict the drug concentration of inhibition and classification models of sensitive/resistance drugs depending on predetermined response levels [10]. Machine learning tools such as the support vector machine [11], random forest [12], neural networks [13], and logistic ridge regression [14] have been utilized for the prediction of drug sensitivity. Riddick *et al.* [15] built an ensemble regression model using random forest to predict in vitro drug response from a signature of gene expression.

In this study, we show that the random forest prediction accuracy significantly improves with prior feature selection and clustering of the data. In addition to the genomic profile of the cell lines, we also incorporate the gene (protein) interaction network and drug (chemical) feature network into the prediction pipeline. There have been a number of different network based studies for drug predictions. For instance, Wang et al. [16, 17] proposed a heterogeneous network model of cell lines, drugs, and targets. Such methods, relying on the matrix multiplication of similarity measures among cell lines and drugs, may make the exclusion (for prediction) of specific drugs difficult (the whole row of the matrix corresponding to that specific drug remains zero). Also, Zhang *et al.* [18] proposed the dual-Layer integrated cell line-drug network model for prediction of drug response. They showed that similar cell lines also respond very similarly to a given drug, and structurally related drugs have similar responses to a given cell line. Following this idea, we show that prior clustering of cell lines and drugs significantly improves the prediction. We compare our results with the Zhang *et al.* extended method in [19] to show that random forest is more effective than the prediction formula provided by [18, 19]. Moreover, in both of these network-based models, the Pearson correlation coefficent has been incorporated to measure the similarity of the samples. Here, we adopt methods from optimal mass transport (OMT) [20–22] to measure the similarity among cell lines and among drugs. To this end, we calculate the Earth Mover’s Distance (EMD) between the probability distributions of given features of the samples in their network of interaction. We showed that Wasserstein distance is more effective than the Pearson correlation in predicting the response. Our drug prediction combines random forest with clustering technique.

In summary, the heterogeneity of pan-cancer cell lines and structurally diverse drugs in large-scale pharmacogenomic databases makes the model fitting for drug prediction difficult. In this work, we present a pipeline for unsupervised clustering of the data in both cell and drug space followed by the prediction of the drug response. Our result shows that clustering of the data significantly improves the prediction accuracy. In addition, clustering helps with feature selection prior to the prediction which makes our algorithm much more efficient. Finally, clustering gives a better insight about the datasets, and facilitates the interpretability of the results and the discovery of key biomarkers.

## Materials and methods

### Data and preprocessing

In this work, we used the anticancer drug response data provided by Drug Sensitivity in Cancer (GDSC) database. GDSC is a publicly available large-scale pharmacogenomic dataset which includes drug screening of more than a thousand human pan-cancer cell lines. The dataset consists of 265 tested compounds including cytotoxic chemotherapeutics as well as targeted therapeutics from various collaborators. GDSC drug responses are in the form of the drug concentration required to inhibit 50% of growth in the proliferation assay (IC50) and the area under the curve for a fitted model (AUC). Genomic profiles of the cell lines within GDSC are obtained and updated directly from the COSMIC cell line project (CCLP) database. In this study we focused on the gene expression data from the cell line project (Affymetrix Human Genome U219 Array). The PPI network data is obtained from the Human Protein Reference Database (HPRD, http://www.hprd.org) [23]. After removing the cell lines that have missing responses for more than 80% of the drugs, we focused on 915 cell lines from 28 tissue types 1.

CCLP and HPRD have about 7,900 genes in common. We further considered making the method computationally more efficient by considering only the 635 genes that are shared in common with OncoKB (Precision Oncology Knowledge Base) database (http://oncokb.org/).

We obtained the drug’s chemical structure from PubChem https://pubchem.ncbi.nlm.nih.gov/. We downloaded the SMILES (Simplified Molecular Input Line Entry Specification) string of 241 drugs for which the PubChem ID is provided in the GDSC database. We also focused on the drugs that had response values for more than half of the cell lines, which ended up yielding 200 drugs. We then extracted the molecular descriptors (drug features) of these drugs from Dragon software (version 7.0) by Kode-Chemoinformatics (https://chm.kode-solutions.net/). The chosen descriptors include properties from the simplest atom types, functional groups, fragment counts, to several properties estimation (such as logP) and drug-like and lead-like alerts (such as the Lipinski’s alert). We initially attained 1,500 molecular descriptors for each drug. This results in a dataset consisting of 183,000 drug-cell line pairs.

### Network-based clustering via Wasserstein distance

Our network-based clustering method is based on the theory of OMT [20–22]. We adopted the 1-Wasserstein distance (EMD) to measure the similarity of the samples in the cell line and drug space. To this end, we assign a probability measure to each sample based on the distribution of its features in their connectivity network. For the cell lines the network of the gene interaction is given by protein-protein interaction (PPI) network. The drug features network is data driven and is built via graphical Lasso. We then utilized the Wasserstein distance to measure the similarity between every pair of probability measures assigned to every two samples (cell lines or drugs). Consequently, we applied these pair-wise distances among samples to perform the hierarchical clustering of the data. In the following sections, first, we will discuss our approach for finding the sample specific probability measures for cells and drugs. We then provide some background of OMT and the Wasserstein distance formulation we used in this work.

### Cell line features in PPI network

Our clustering of cell lines is based on their gene expression in the PPI network. We constructed a weighted graph by considering the given gene interaction network as a Markov chain [24, 25]. Consider a gene *i* and its neighbor genes *j ∈ N* (*i*) in their interaction network (here in HPRD) for a given sample. Let *ge*_*k*_ stand for the expression level of gene *k* in a given sample. The principle of mass action allows us to compute the probability of the interaction of gene *i* to gene *j* (*p*_*ij*_) to be proportional to their expression, i.e. *p*_*ij*_ ∝ (*ge*_*i*_)(*ge*_*j*_) [26]. By normalizing *p*_*ij*_ such that such that Σ_*j*_*p*_*ij*_ = 1, we have the stochastic matrix *p* of the Markov chain associated to the network as follows:

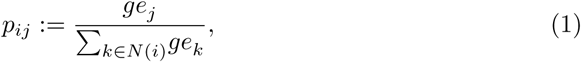

The Markov chain given by (1) reaches a stationary distribution which is invariant under a right multiplication by *p*, i.e. *πp* = *π*, [24]. Solving this formula for the special stochastic matrix *p* defined by 1, *π* has the explicit expression:

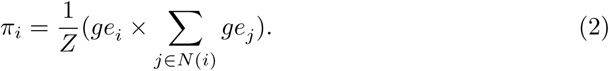

*Z* is a normalization factor forcing *π* to be a probability vector. Of note, this normalization is necessary since we want to consider the invariant measure to be a probability distribution over all genes for each specific sample. The invariant measure defined by 2, gives a value to each gene which is not only dependent on the gene expression of the gene *i*, but also on the total gene expressions of the neighboring genes *j ∈N* (*i*). For each sample, we calculated a vector *π* = (*π*_*i*_)_*i*=1,…,*n*_ for all the *n* genes in our data sets. We then calculate the EMD to find the distance between a pair of vectors of the form *π* assigned to every two samples. In fact, we measured the similarity between samples by finding the EMD between the distributions of the invariant measures assigned to the cell lines. As we discuss later 5, the EMD utilized both the invariant measures and PPI interaction network connectivity to measure the similarity between cell lines.

### Network of drug features via graphical lasso

We initially extract molecular descriptors of the 200 drugs from Dragon software. The following descriptors have been removed: 3D based descriptors (since the 3D information is not provided), descriptors that are constant or near constant, and descriptors with missing values. This results in 1,500 drug features. We then remove the highly correlated drug features via unsupervised clustering of the features. We perform the hierarchical clustering of the features using the Spearman’s rank-order correlation between features for all 200 drugs 2(a). We then choose a representative feature of each cluster randomly. This reduced the number of features to 500 that are not significantly correlated. We then build a network of these drug features which will be used later for clustering of the drugs via Wasserstein distance. We make the connected network of the 500 features sparse via graphical lasso (GL) so that obtaining the Wasserstein distance would be more computationally efficient.

In an undirected graph, each vertex can represent a random variable. Now, consider *n* observations have a multivariate Gaussian distribution with mean *µ* (=0) and covariance matrix Σ. The GL [27] is a regularization framework for estimating the covariance matrix Σ, under the assumption that its inverse (precision matrix), Θ = Σ^−1^, is sparse [28]. If an element *θ*_*jk*_ = 0, this implies that the corresponding variables (vertices) of indices *j* and *k* are conditionally independent, given other variables. This can justify removing the edge connecting these two vertices (*j* and *k*). GL impose as *ℓ*_1_ penalty for estimation of Σ to increase this graph sparsity. The GL problem minimizes a *ℓ*_1_-regularized negative log-likelihood as follows:

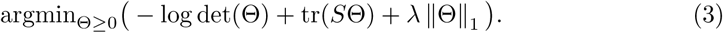

where *S* is the empirical covariance matrix, ‖Θ‖_1_ denotes the sum of the absolute values of Θ, and *λ* is a tuning parameter controlling the amount of *ℓ*_1_ shrinkage. Here, we tune *ℓ*_1_ to make the network as sparse as possible so long that it does not decrease the accuracy of drug response prediction. We then consider for each specific drug a weighted network by assigning the drug’s feature values to the nodes of the network obtained from GL. As before, we further normalize these value to sum up to one to be considered a probability distribution. Here, we do not apply the integrative measure used in the gene interaction network case, since the data driven network of drug features here is representing the correlation between the nodes (drug feature) rather than the physical interaction among them. After assigning the probability distributions to these drugs, we calculate the EMD to find the similarity measure among the drugs. These similarity measures are based on both feature values of the drug chemical structures and the network connectivity of these features 5.

### Clustering via Wasserstein distance

We use the theory of OMT to define distances among samples. OMT is a rapidly developing area of research that deals with the geometry of probability densities [20]. The subject began with the work of Gaspard Monge in 1781 [29] who formulated the problem of finding minimal transportation cost to move a pile of soil (“deblais”), with mass density *ρ*^0^, to an excavation (“remblais”), with a mass density *ρ*^1^. A relaxed version of the problem was introduced by Leonid Kantorovich in 1942 [30]. We will only give a special case of the general problem that we will need in the present work. This analysis method is appropriate when location, in this case represented by a network, reflects relatedness and correlation. Thus, moving signal between neighboring nodes results in a small contribution to the distance measure, whereas different activity levels in distant neighborhoods of the network contribute significantly.

Let *ρ*^0^, *ρ*^1^ ∈ *P*(Ω) where Ω ⊆ ℝ^*N*^ and 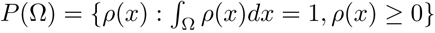 The 1-Wasserstein distance, also known as the Earth Mover’s Distance (EMD), is defined as follows:

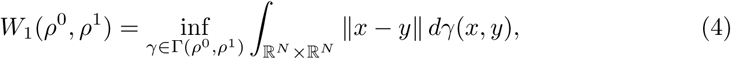

where Γ(*ρ*^0^, *ρ*^1^) denotes the set of all couplings between *ρ*^0^ and *ρ*^1^, that is the set of all joint probability measures *γ* on Ω *×* Ω whose marginals are *ρ*^0^ and *ρ*^1^. Here, the cost function of the transportation is defined as the ground distance *d*(*x, y*) = ‖*x* −*y*‖.

The optimization problem (4) has an analogous formulation on a weighted graph. Let us consider a connected undirected graph *𝒢* = (*𝒱, ε*) with *n* nodes in *V* and *m* edges in *ε*. Given two probability densities *ρ*^0^, *ρ*^1^ *∈* ℝ *n* on the graph, the EMD problem seeks a joint distribution *ρ* ∈ ℝ *n*×*n* with marginals *ρ*^0^ and *ρ*^1^ minimizing the total cost ∑*c*_*ij*_*ρ*_*ij*_:

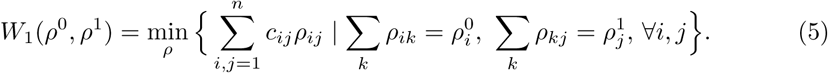

Here *c*_*ij*_ is the cost of moving unit mass from node *i* to node *j* and is taken to be the shortest path to go from node *i* to node *j*. For implementations, we use the Hungarian algorithm [31] to compute the EMD on our reference networks. We compute the pairwise EMD of samples for unsupervised clustering of the GDSC cell lines and drug data. Each sample (cell or drug) is represented as a vector of its pairwise distances to all other samples (its distance to itself is zero). We then apply the hierarchical agglomerative clustering to the sample vectors. Hierarchical clustering imposes a hierarchical structure on the samples and their stepwise clusters. To achieve a certain number of clusters, the hierarchy is cutoff at the relevant depth. Here, we choose the optimal number of clusters based on the silhouette values [32]. The silhouette value for each sample is a measure of how similar that sample is to other samples in its own cluster compared to samples in other clusters. The *silhouette value* for the sample *i, s*(*i*) is defined as:

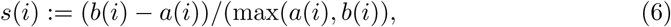

where *a*(*i*) denotes the average distance of the sample *i* to all samples within its own cluster (squared Euclidean distance between sample vectors), and *b*(*i*) denotes the minimum average distance of the sample *i* to samples of other clusters. Silhouette values range from −1 to +1. A high silhouette value indicates that the data are appropriately clustered. Therefore, we choose the optimal number of clusters by analyzing the average silhouette values of samples to make sure they stay close to 1.

### Prediction of drug response in paired cell line-drug clusters

We apply random forest for predicting drug response in each paired cluster of cell lines and drugs. Random forest is an efficient ensemble learning algorithm capable of handling high-dimensional data. Here, we use random forest for supervised learning of a regression problem. We choose the number of decision trees to be 100 and the number of randomly selected features to be *N/*3 where *N* is the number of features as recommended for regression problems in [33]. Increasing the number of decision trees or selected features make the algorithm computationally expensive; however, it does not improve the performance (prediction) significantly. We also compare our result to the network-based model introduced in [19] (CDCN model) which is the extension of Zhang’s dual layer integrated cell line-drug network model [18]. The integrated model does not consider the clustering of the data, but it uses the similarity measure between cell lines and drugs via Pearson correlation. Our method can apply the closed formula proposed in these works instead of random forest in the prediction pipeline. Assume that *r*(*c, d*) is the IC50 value of the pair of drug *d* ∈ 𝒟 and cell line *c* ∈ 𝒞 where 𝒟 and 𝒞 denotes a set of drug or cell lines correspondingly. For a new cell line *c*^*^ and a new drug *d*^*^, we would like to predict the drug response, *r*(*c*^*^, *d*^*^), based on the known values *r*(*c, d*). Using the metric *d*_𝒟_ (EMD), we can cluster 𝒟 ∪ {*d*^*^}. Denote by 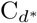 the cluster in which *d*^*^ lies, but with *d*^*^ removed from it. Similarly, we can compute 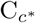 in the cell line space using the *d*_*𝒞*_. We define a similarity weight function as 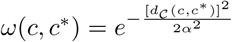 between cell lines and the similarity weight function between drugs defined as 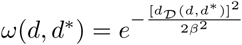 with the vector of decay parameters, i.e. *ζ* = (*α,β*). These similarity weights are defined in such a way that their values are higher when the samples are more similar to each other. Therefore, in case of using Pearson correlation for these similarity measures, we substitute *d*_*𝒞*_ with 1 − *ρ*_*𝒞*_ for cell lines and *d*_*𝒟*_ with 1− *ρ*_*𝒟*_ for drugs, where *ρ* denotes the Pearson correlation.

Consequently, we compute the drug response by:

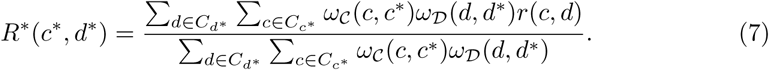

The decay parameter *ζ* = (*α,β*) can be optimized on the training set Γ by minimizing the error of response prediction as follows [19]:

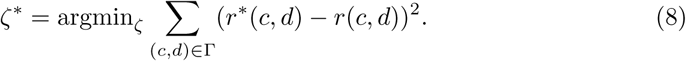

where *r*^*^(*c, d*) is the prediction of drug response *r*(*c, d*) in the training set.

## Results

Our prediction pipeline starts with unsupervised clustering of cell lines and drugs. The hierarchical clustering of the cell lines results in 6 clusters with optimal silhouette mean values. Figure 3 illustrates the results of clustering for cancer types with more than 20 cell lines. As shown in 3, cluster 1 perfectly separates the liquid cancers of leukemia and lymphoma and almost nothing from the solid tumors. This could significantly improve the prediction given liquid tumors responded very differently to the anti cancer agents than solid neoplasms [34].

**Fig 1.**
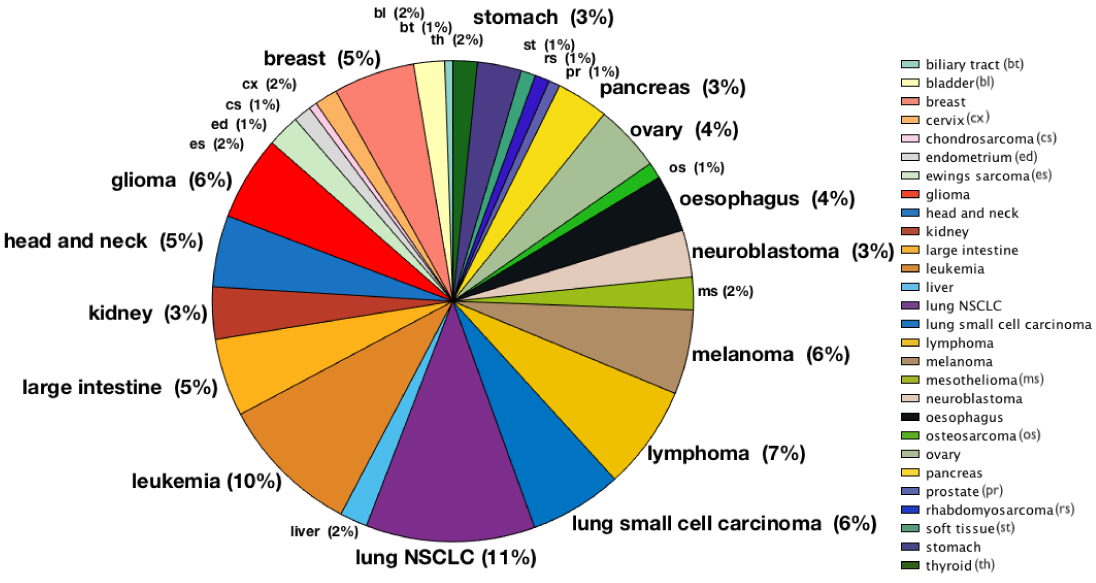
Distribution of the 915 cell lines from GDSC with respect to their tissue types. The largest cancer types are lung NSCLC (11 %) and leukemia (10%). Lymphoma (7%), glioma (6%), melanoma (6%) and lung small cell carcinoma (6%) are the next major types.

**Fig 2.**
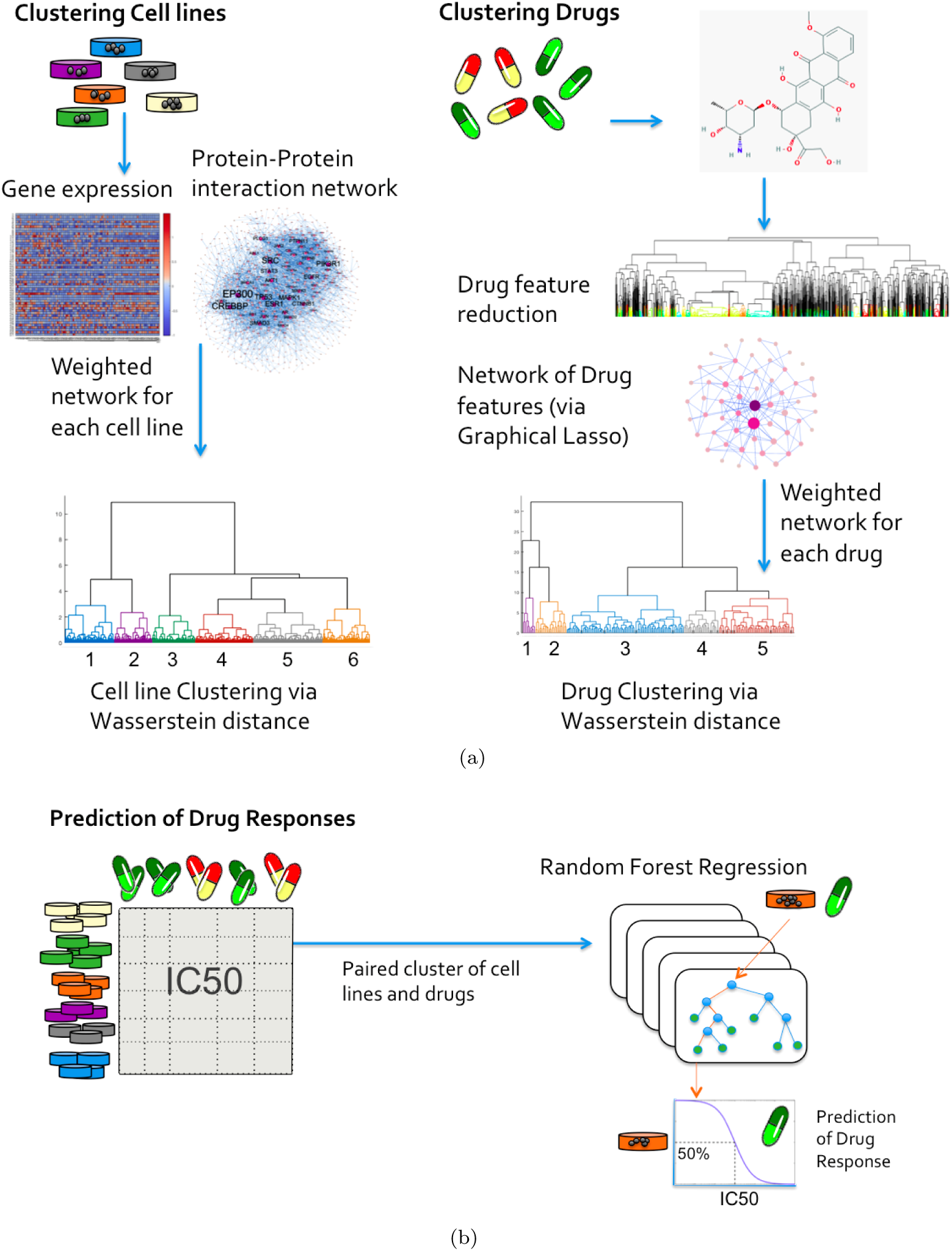
Overview of network-based clustering and prediction of drug response. (a) Clustering of cell lines and drugs. The gene expression profile is provided by GDSC for 915 cell lines and the gene (protein) interaction network is obtained from HPRD. Our clustering algorithm considers the expression of a specific gene in its neighborhood interaction network and utilizes EMD to measure the similarity between cell lines. For the drugs, we first obtain the chemical features from Dragon software. We then use hierarchical clustering to remove the highly correlated features. Next, we build a data driven network of chemical features using graphical lasso. Again, we utilize EMD to perform the hierarchical clustering of the drugs. (b) Prediction of drug responses. We predict the drug response in each paired cluster of the cell lines and drugs which are small dashed blocks in the matrix of IC50 values in the figure. We perform the 3-fold cross validation by applying the random forest regression to each paired cluster of cell lines and drugs. Random forest model can be replaced by the closed formula presented in the “Prediction of drug response in paired cell line-drug clusters” section.

**Fig 3.**
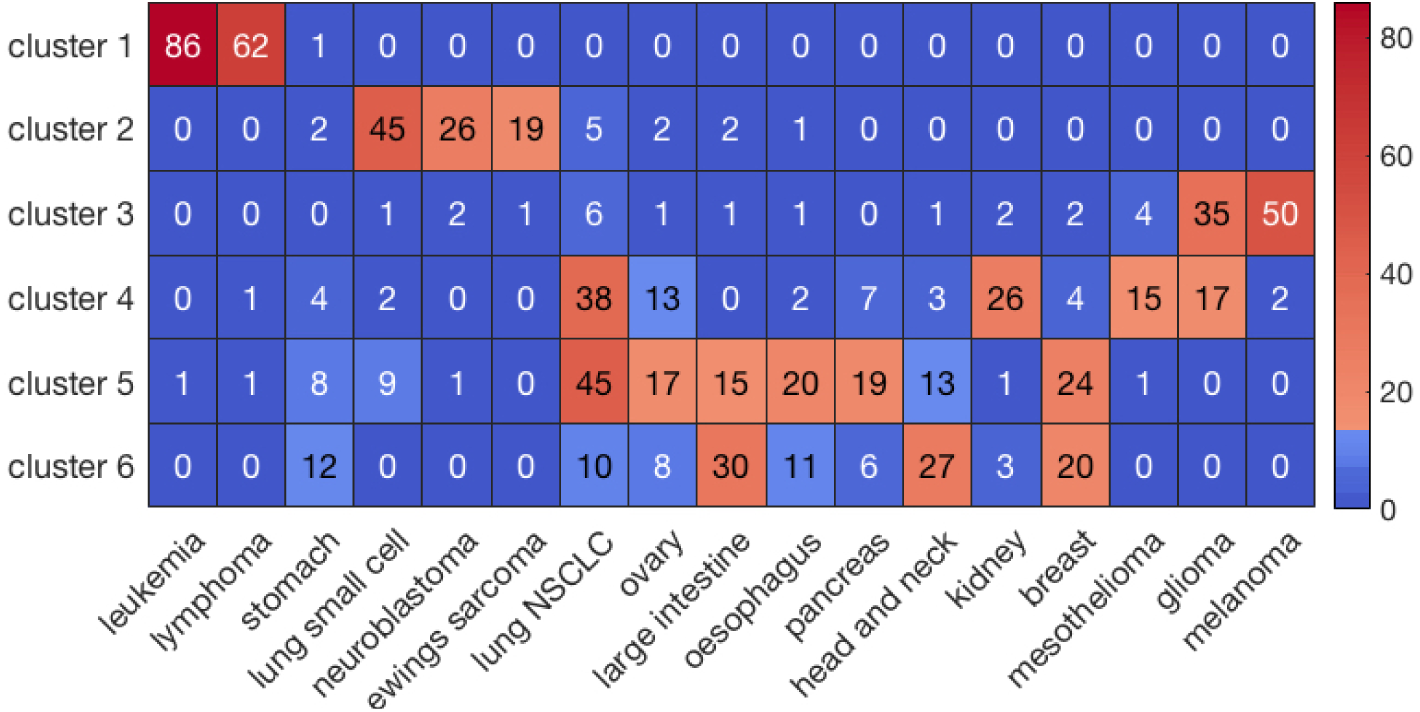
The majority of cell lines within a specific cancer type have been clustered together including leukemia, lymphoma, lung small cell, neuroblastoma, ewing sarcoma, oesophagus, pancreas, mesothelioma, and melanoma. Lung NSCL and breast cancer have been distributed between two neighboring clusters. The neighboring clusters 4,5, and 6 are close to each other and are all members of a bigger cluster in the hierarchical clustering results 2(a). These clusters are more heterogenous than clusters 1, 2, and 3.

On the other hand, hierarchical clustering of the drug results in 5 clusters. Consequently, for each of the 30 paired clusters of cell line and drugs we trained a random forest. In each paired cluster, we performed the 3-fold cross validation, i.e., we used 67% of the drug responses as the training set to build the random forest model, then we applied these models to predict the drug response in the remaining 33% of the data. After performing the 3-fold cross validation in each paired cluster, we concatenated the prediction for all the pairs of cell and drugs in the paired cluster. Figure 4 provides box-plots of distribution of correlations and coefficient of determination of the predicted and observed IC50’s in the 30 paired cluster of cell lines and drugs. To gain the accuracy of prediction for the whole data set, we concatenated the prediction of all clusters and then calculate the correlation and coefficient of determination of the predicted and observed values 1. Also, taking the average value of the correlations and coefficient of determinations of the clusters results in very similar values 4. We also performed a 3-fold cross validation via random forest without prior clustering of the data. We trained the random forest using the initial features of cell lines and drugs (635 genes and 500 molecular descriptors) using 67% of the dataset at each fold. Similarly, we concatenated the results to have a prediction for all the pairs in our dataset. As we see in Table 1, the prior clustering of the data significantly improves the prediction accuracy. Among our 6 clusters of cell lines, cluster 1 and 3 have the best prediction accuracy. These two clusters are also the least heterogenous in terms of the cancer types which can explain their improved prediction. This also supports the fact that reducing the complexity of data with prior clustering improves the prediction accuracy. Figure 5 shows the correlation and coefficient of determination in the best and worst paired cluster (among 30 paired clusters) with regards to the prediction accuracy. The best prediction belongs to the pair of third and first clusters of the cell line and drug, respectively 2(a). This cluster of cell lines mainly consists of glioma and melanoma 3.

**Table 1.** Performance comparison of four models for 183,000 pairs of cell line-drugs in GDSC based on correlations (R) and coefficient of determination (R^2^) of the predicted and observed IC50’s. Applying EMD for prior clustering of data improves the performance of random forest. Also, the integrated model via EMD has a better performance than the integrated model using Pearson correlation.

| Model | $R$ | $R^2$ |
| --- | --- | --- |
| Random Forest w/ prior EMD Clustering | 0.89 | 0.79 |
| Integrated (w/ EMD) | 0.86 | 0.59 |
| Random Forest | 0.77 | 0.60 |
| Integrated (w/ Pearson correlation) | 0.74 | 0.53 |

**Fig 4.**
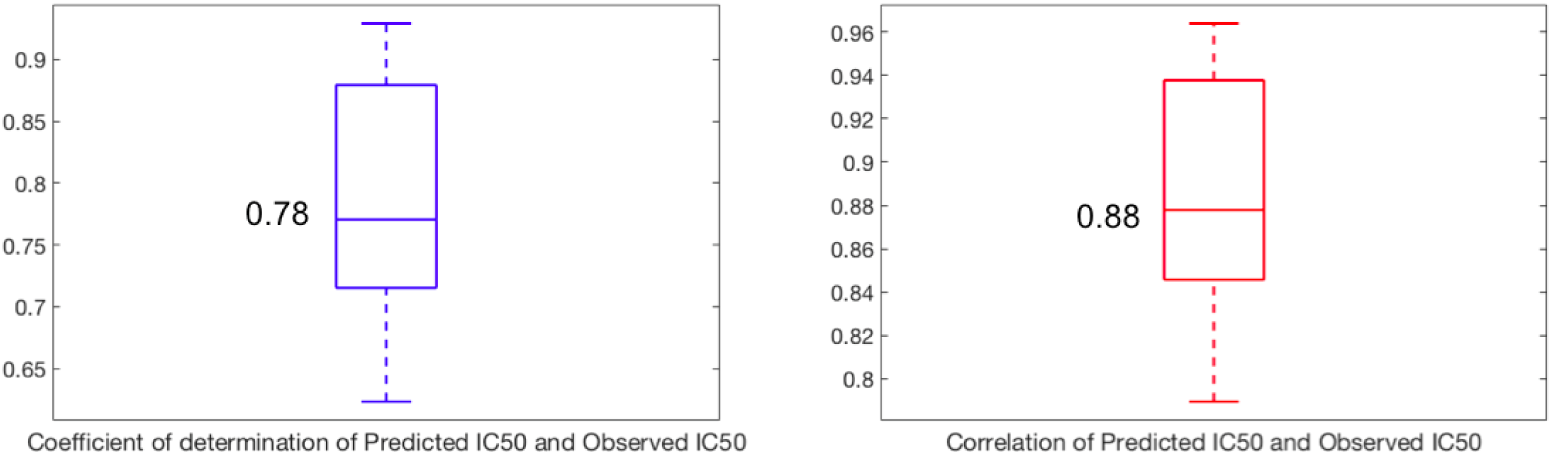
The distribution of correlations (R) and coefficient of determination (R^2^) of the predicted and observed IC50’s in the 30 paired clusters of cell lines and drugs. The average values of R and R^2^ which are written in the plots show the high accuracy of prediction within the clusters.

**Fig 5.**
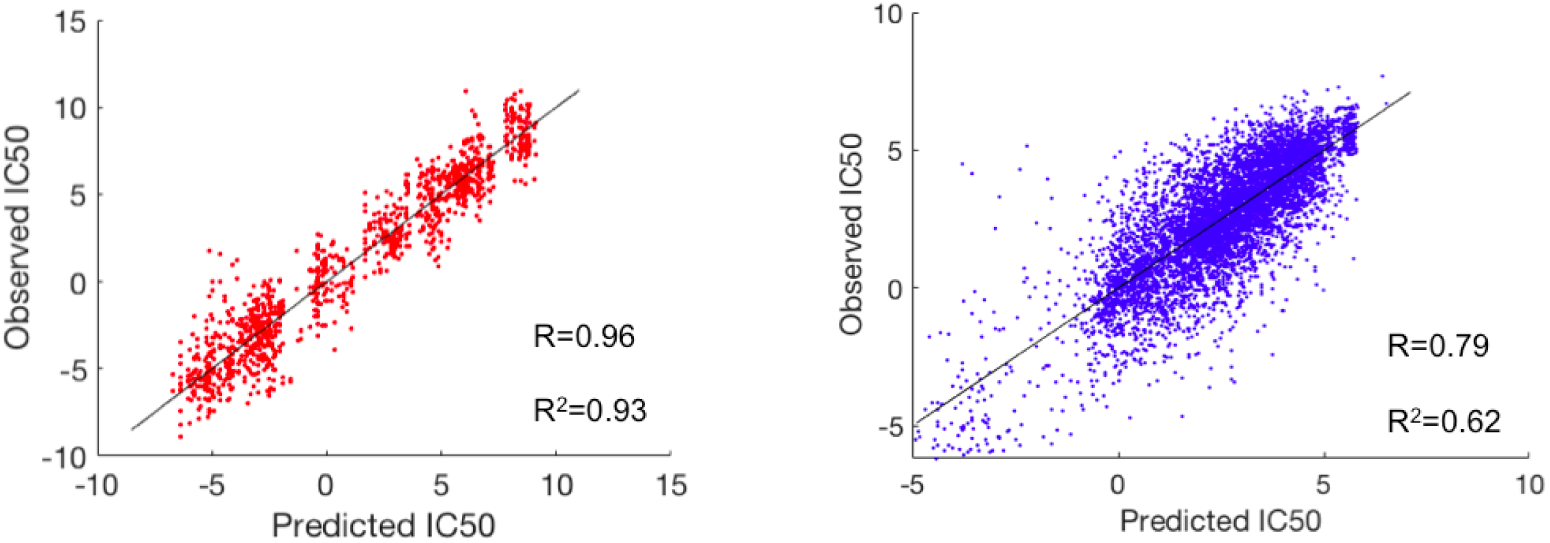
Best (red) and worst (blue) of 30 clusters with respect to the prediction accuracy. The best prediction belongs to the pair of third cluster of the cell lines (mainly glioma and melanoma) and first cluster of drugs, and the worst prediction belongs to the sixth cluster of cell lines (mainly consisting of breast, head and neck, large intestine, and stomach cancer) and fifth cluster of drugs 2(a).

Our prediction pipeline is also interpretable and can be used to discover molecular biomarkers. Here, we obtained the significant genes in this paired cluster that are contributing the most in both prediction and clustering of the cell lines. To this end, we first focused on the paired clusters of cell lines and drugs that had the best drug response (lowest average IC50) which belonged to first cluster of both cell lines and drugs 2(a). We derived the feature importance score from the random forests algorithm corresponding to the prediction in this cluster as suggested in [33]. This provided ranking of the genes based on their contribution in the construction of decision trees within the random forest model that is used for the response prediction. We also investigated genes that are differentially expressed for this cluster of cell lines compared to the rest of the cell lines by performing the t-test. Among the top 200 genes (from the random forest feature importance score), we selected 70 genes based on their Bonferroni corrected p-values from the t-test (p-value *<* 0.05). These significant genes are important features for both the drug response prediction and clustering of the cell lines (in cluster 1). The PPI network of these genes was constructed via MetaCore software and is presented in Figure 6.

**Fig 6.**
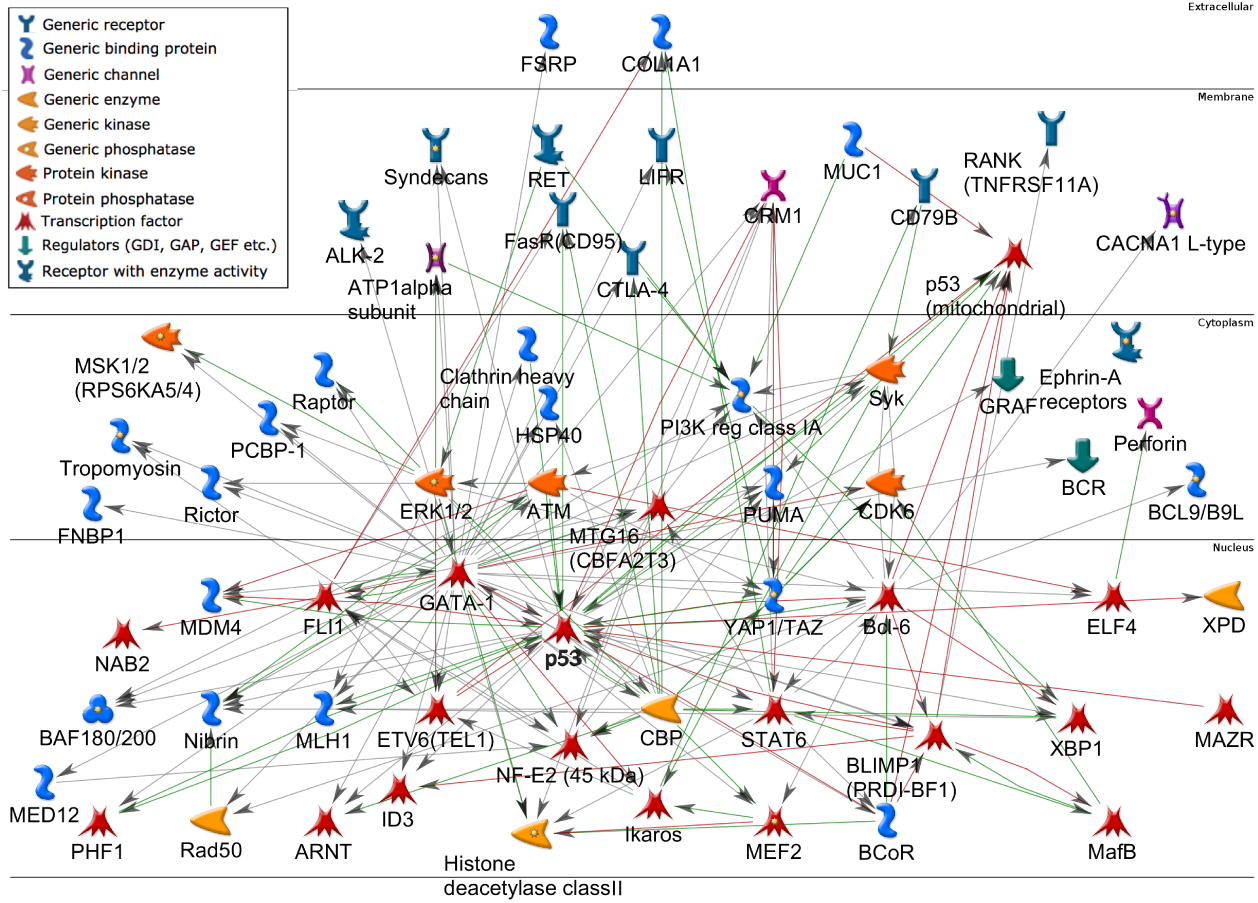
PPI network of significant genes for the paired cluster of cell line and drug with best drug response. The genes are identified based on their feature importance from random forest prediction and their differential expression (using t-test) between this cluster and the remaining cell lines.

## Discussion

In this study, we developed a framework for clustering and predicting drug sensitivity of pan cancer cell lines in the GDSC database. Our pipeline consists of three steps: unsupervised clustering of the cell lines, unsupervised clustering of the drugs, and prediction of drug response in paired cluster of cell lines and drugs. We studied the responses of a cohort of 183,000 drug-cell line pairs from the GDSC dataset. The novelty of our method is in prior clustering of the data (both cell lines and drugs) by utilizing EMD from the theory of optimal mass transport. It has been shown that similar cell lines by their gene expression profiles exhibit similar responses to the (structurally) similar drugs [18, 19]. We improved these methods by focusing on the paired cluster of cell line and drugs to which a new pair of cell line-drug in the test set belongs. The similarity measures obtained from the EMD improves the prediction accuracy of drug responses compared to Pearson correlation in the integrated model [18]. We also performed the prediction using the random forest model, which results in the better coefficient of determination of the predicted and observed responses as well as a better correlation 1. Previous work has shown that pre-processing data by a clustering algorithm improves prediction accuracy of even random forest models [35]. Here, we also observed that applying random forest with prior clustering has better prediction performance than using random forest without the clustering 1. In fact, the best prediction accuracy belonged to the cell line clusters that were less heterogeneous (cluster 1 and 3 3).

In many biological applications of machine learning techniques, it is equally important to not only have an accurate prediction, but also an interpretable model. We showed that we can derive significant genes of both clustering and prediction by performing the t-test and using the importance score from random forest. We analyzed the cluster with the best drug response (lowest average IC50). This cluster consisted of mostly leukemia and lymphoma cell lines. Furthermore, drugs in this cluster mainly target mitosis and DNA-replication including antimetabolites. The PPI network of these genes is shown in Figure 6.

The proxy of cancer patients (in vivo) with cell lines (in vitro) to characterize the response against therapeutic agents needs to be fully investigated. However, such proximation could provide candidate therapeutic agents to help with the systematic study of drug responses on a large number of patients which is costly, time-consuming and limited to the scope of drugs that have been approved and tested on patients. On the other hand, the identification of a drug sensitivity biomarker is essential for precision medicine. This study illustrates a novel mathematical network-based model for prediction of cell line-drug responses as well as identification of relevant genomic biomarkers and biological processes for anti-cancer drug treatment.

## Acknowledgments

This study was supported by AFOSR grant (FA9550-17-1-0435), NIA grant (R01-AG048769), MSK Cancer Center Support Grant/Core Grant (P30 CA008748), and a grant from Breast Cancer Research Foundation (grant BCRF-17-193).

